# POCKET-seq enables genome-wide profiling of on- and off-target transcriptional regulation events by dCas9-KRAB during CRISPR interference experiments

**DOI:** 10.64898/2026.06.01.729390

**Authors:** Christine M. Joyce, Griffin D. Kramer, Jonathan T. Vu, Katherine U. Tavasoli, Brooke M. Gardner, Chris D. Richardson

## Abstract

CRISPR interference screens use catalytically inactive dCas9 fused to a repressor domain to enable genetic perturbations at the transcriptomic level. Interpretation of results involves identification of guide RNAs associated with the screen phenotype, followed by secondary analysis. During validation of a genetic screen, we observed different phenotypes from non-overlapping guide RNAs targeting one gene. Here, we developed POCKET-seq to map the binding of dCas9 genome-wide. We show that off-target binding occurs frequently and can generate false-positive interactions when it occurs near the promoter of genes associated with the screen phenotype. POCKET-seq classifies these false-positive and true-positive interactions using gene ontology.

## Background

Genetic screens enable researchers to identify genes involved in a biological pathway, process, or phenotype. In human cell lines, CRISPR interference (CRISPRi) is a common tool for forward genetic screens [1–3]. One implementation of pooled CRISPRi screens uses a guide RNA (gRNA) library and a catalytically dead Cas9 (dCas9) fused to a transcriptional repressor domain, such as Krüppel associated box (KRAB) [3]. This fusion protein can therefore be targeted to the promoter regions of genes and in turn downregulate gene expression [1,2].

CRISPRi screens work by tracking the abundance of different gRNAs during selective pressure. Depleted gRNAs repress genes that would otherwise promote survival, while enriched gRNAs repress genes that would otherwise sensitize the cells. These differences in gRNA abundance are uncovered by sequencing the gRNAs present at the end of the experiment and mapping them back to a reference dataset, producing a count table for every gRNA in the library [4,5]. The gRNA counts are then converted to phenotype and significance scores and rank ordered, and the top hits are chosen for validation in orthogonal assays.

Despite the ubiquity of CRISPRi screens, there is no established methodology for managing variability. As validation experiments require considerable resources, inconsistent results can incur costly delays. During analysis of a genetic screen, we discovered two non-overlapping gRNAs produced vastly different phenotypes [6]. This observation was consistent with an off-target binding event of dCas9-KRAB that regulated an unknown gene, which in turn affected our phenotype.

To define the nature of these binding events, we developed a genomic technique to identify off-target interactions in transcriptional regulation experiments, which we have termed <u>P</u>rofiling <u>O</u>ff-target <u>C</u>RISPR <u>K</u>nockdown through <u>E</u>pitope <u>T</u>ag (POCKET-seq). Using ChIP-seq of HA-tagged dCas9-KRAB, POCKET-seq reveals off-target binding interactions and classifies these interactions according to their likelihood to produce false-positive or true-positive outcomes.

## Results and Discussion

We performed a genome-wide CRISPRi screen to identify genes regulating peroxisome import in HCT116 human colorectal cancer cells [6]. The screen relied on sequestration of a zeocin resistance construct (mVenus-ZeoR-PTS) within peroxisomes. In this approach, cells with active peroxisome import machinery sequester the zeocin resistance (ZeoR) construct within their peroxisomes, making them sensitive to zeocin. Conversely, cells lacking functional import retain the construct in the cytosol, rendering them resistant to zeocin. Candidate genes from the primary screen were then individually re-inhibited with CRISPRi using two different gRNAs per gene, and peroxisome abundance was monitored using fluorescence microscopy for mVenus-PTS foci and immunofluorescence staining of the peroxisomal membrane protein PMP70.

Individual gRNAs targeting the gene *Suppressor of Cytokine Signaling 3 (SOCS3)* produced contradictory phenotypes in our secondary analysis: *SOCS3* guide 1 (SOCS3_g1) decreased peroxisome abundance, while *SOCS3* guide 4 (SOCS3_g4) had no effect compared to non-targeting controls (NTCs) (**Fig. 1A, Additional file 1: Fig. S1A-C**). Although RT-qPCR confirmed both gRNAs depleted *SOCS3* (**Additional file 1: Fig. S1D**), lentiviral re-expression of *SOCS3* cDNA failed to restore peroxisome abundance in the presence of SOCS3_g1 (**Additional file 1: Fig. S1E**). Taken together, this data suggested the SOCS3_g1 phenotype was driven by genuine, off-target regulatory interactions.

**Figure 1.**
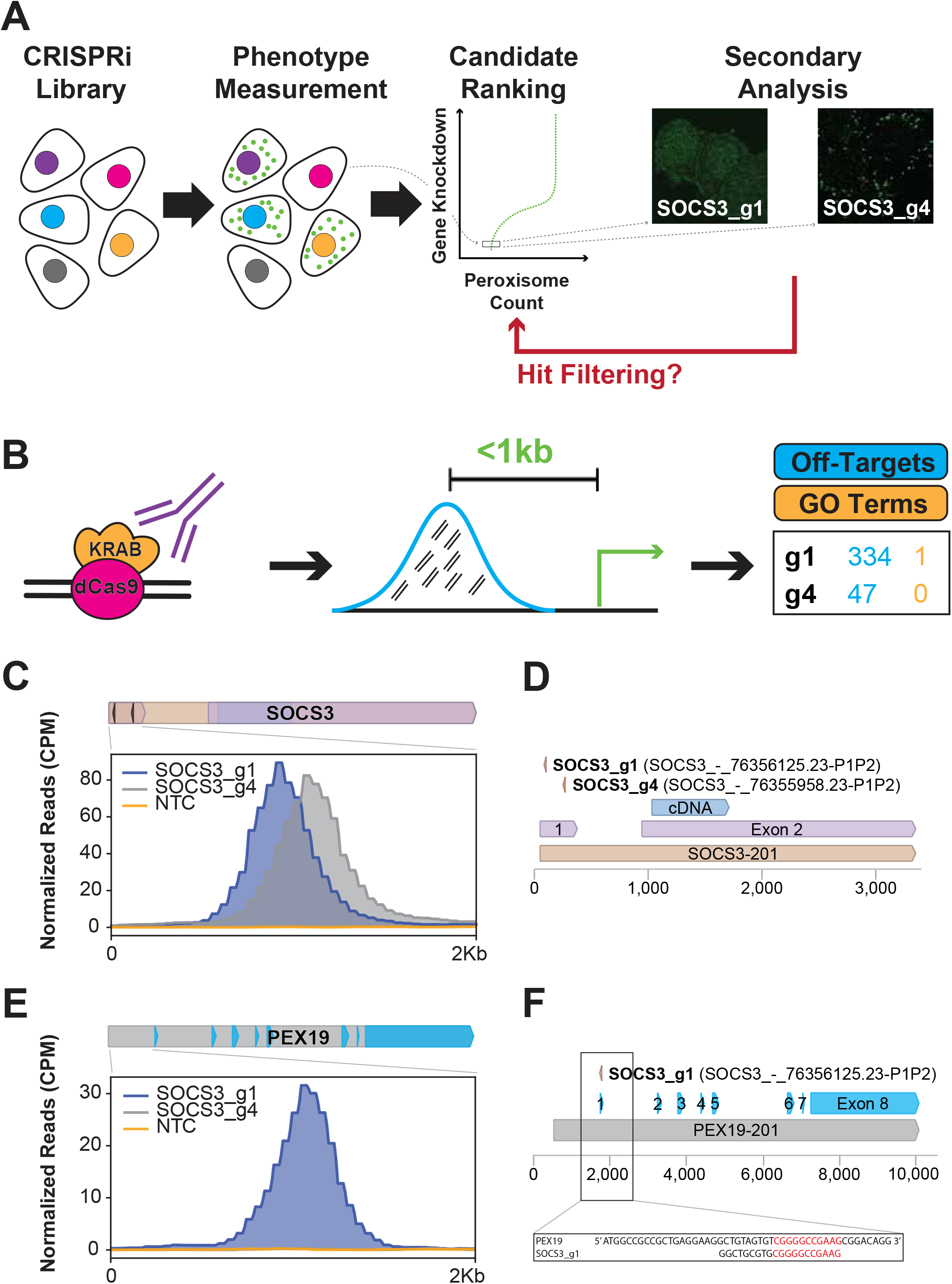
CRISPR interference screens can result in off-target transcriptional repression events. A. Schematic of CRISPRi screen workflow. A library of genome wide CRISPRi cells were generated and monitored for peroxisome abundance. gRNAs targeting specific genes were ranked according to the magnitude of peroxisome depletion. Secondary analysis using microscopy for a fluorescent reporter (mVenus) with a peroxisomal targeting signal (PTS) revealed that one *SOCS3* gRNA depleted peroxisomes, while the other did not. B. Schematic of POCKET-seq workflow. Chromatin immunoprecipitation (ChIP) was performed for the HA epitope tag on dCas9-KRAB. Reads were aligned to the reference genome and transcription start sites (TSS) within 1kb were identified. Off-targets for SOCS3_g1 and SOCS3_g4 were quantified and membership within the gene ontology (GO) term “peroxisome organization” was assessed. C. Normalized ChIP-seq read counts quantifying the binding of dCas9-KRAB to the on-target site, which is the promoter region of the *SOCS3* gene. Immunoprecipitations were performed using an HA-tag antibody from CRISPRi non-targeting control (NTC) and *SOCS3*-depleted (SOCS3_g1 and SOCS3_g4) HCT116 cells. All plots are representative of *n* = 2 biological replicates. D. Schematic of the gRNAs for the *SOCS3* gene. E. Normalized ChIP-seq read counts quantifying the binding of dCas9-KRAB to the off-target site, which is the promoter region of the *PEX19* gene. Immunoprecipitations were performed using an HA-tag antibody from CRISPRi non-targeting control (NTC) and *SOCS3*-depleted (SOCS3_g1 and SOCS3_g4) HCT116 cells. All plots are representative of *n* = 2 biological replicates. F. Schematic of the partial overlap of SOCS3_g1 for the *PEX19* gene.

To map these on- and off-target binding sites, we developed POCKET-seq, which consists of chromatin immunoprecipitation of the HA epitope tag on dCas9-KRAB (**Fig. 1B, Additional file 2: Fig. S2A**). Bioinformatic processing of this data maps ChIP-seq reads to the reference genome, identifies transcription start sites (TSS) located within 1kb, and ranks binding events by magnitude of ChIP signal, gene ontology (GO) term membership, or primary screen phenotype scores. Our pipeline highlights off-target interactions most likely to produce the observed false-positive phenotypes.

Applying POCKET-seq revealed 334 off-target dCas9-KRAB interactions for SOCS3_g1 and 47 for SOCS3_g4 (**Fig. 1B**). While both guides properly loaded dCas9-KRAB to the *SOCS3* promoter as intended (**Fig. 1C-D**), only SOCS3_g1 had an off-target site within a gene in the “peroxisome organization” GO group; conversely, none of the 47 off-targets for SOCS3_g4 fell into this category (**Fig. 1B**). Specifically, SOCS3_g1 targeted *PEX19*, a critical peroxisomal regulator (**Fig. 1E-F**). RT-qPCR confirmed SOCS3_g1 directly depleted *PEX19* transcripts, whereas SOCS3_g4 did not (**Additional file 2: Fig. S2B**). Furthermore, adding back *PEX19* cDNA rescued the peroxisome abundance phenotype, confirming the screen result was dependent on the direct, off-target inhibition of *PEX19* by SOCS3_g1 (**Additional file 2: Fig. S2C-D**).

To validate the accuracy of POCKET-seq for mapping dCas9-KRAB transcriptional regulation events, we cross-validated our approach with other nuclease-mapping approaches using gRNAs from this study as well as published gRNAs targeting VEGFA site 2 and HEK293 site 4 (**Fig. 2A**). We found almost no overlap with *in silico* predictions; the CRISPOR tool [7] identified 191 off-targets for SOCS3_g1 but did not predict the critical *PEX19* interaction (**Fig. 2B-C**). Because CRISPOR relies primarily on sequence mismatch rules, we propose it is unable to account for the local chromatin context of potential binding events, leading to low overlap with POCKET-seq.

**Figure 2.**
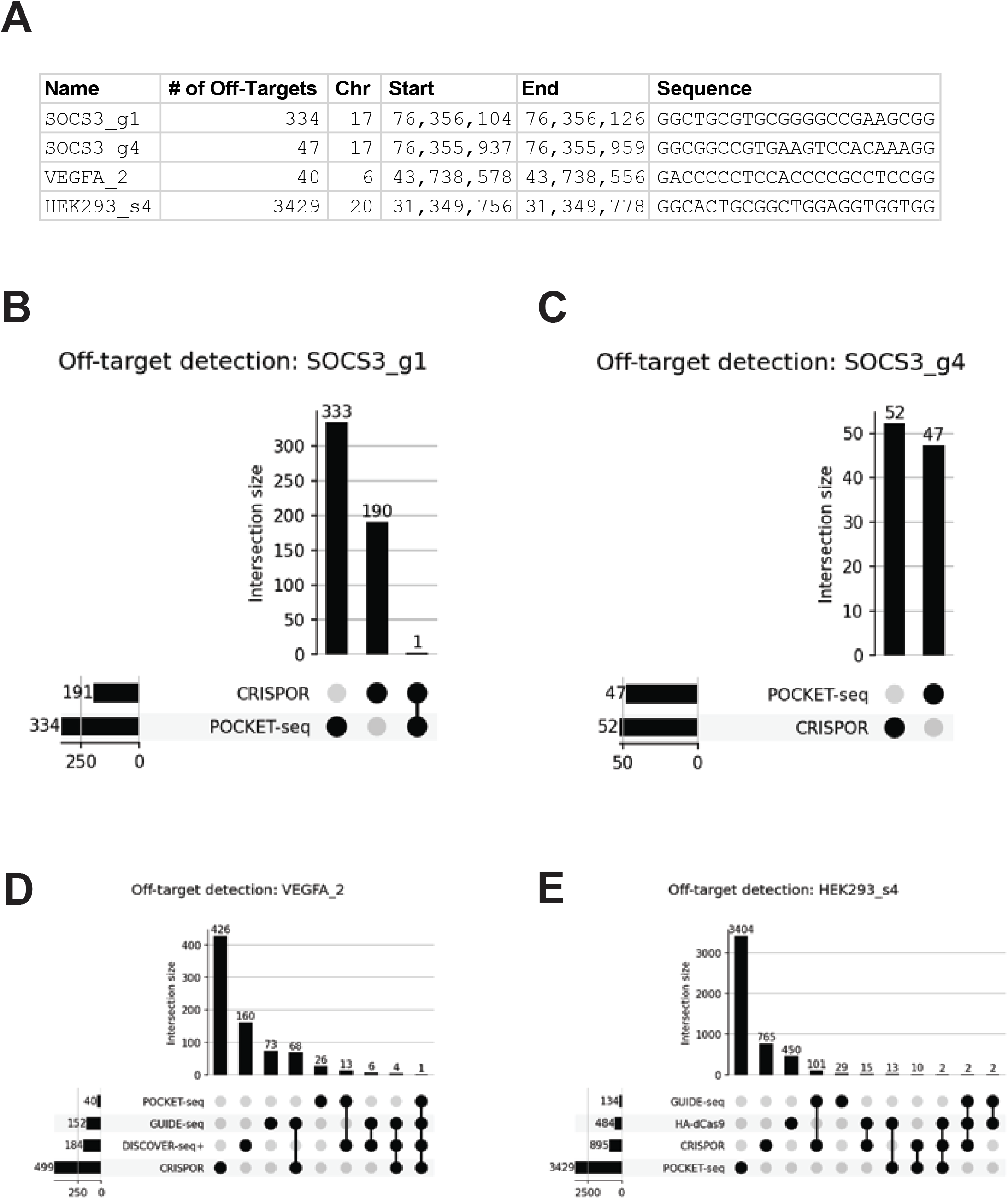
ChIP-seq of dCas9-KRAB can be used to identify off-targets of CRISPR interference. A. Table of genomic chromosomal locations, guide RNA sequences, and number of off-target dCas9-KRAB binding sites identified by POCKET-seq for each of the four guide RNAs examined in this study (SOCS3_g1, SOCS3_g4, VEGFA_2, and HEK293_s4). B. UpSet plots comparing overlap between off-target sites predicted *in silico* by CRISPOR [7] and measured by POCKET-seq in HCT116 cells for SOCS3_g1. Of the 334 sites identified by POCKET-seq, and the 191 sites predicted by CRISPOR, there was only 1 overlapping off-target site, which did not correspond to the relevant *PEX19* off-target. C. UpSet plots comparing overlap between off-target sites predicted *in silico* by CRISPOR [7] and measured by POCKET-seq in HCT116 cells for SOCS3_g4. Of the 47 sites identified by POCKET-seq, and the 52 sites predicted by CRISPOR, there were no overlapping off-target sites. D. UpSet plots comparing overlap between off-target sites predicted *in silico* by CRISPOR[7] and measured by GUIDE-seq in U2OS cells [8], DISCOVER-seq+ in HEK293T cells [10], and POCKET-seq in HCT116 cells for VEGFA_2. CRISPOR predicted 499 sites. GUIDE-seq identified 152 sites. DISCOVER-seq+ identified 184 sites. POCKET-seq identified 40 sites. Overlaps are quantified between the various techniques. E. UpSet plots comparing overlap between off-target sites predicted *in silico* by CRISPOR [7] and measured by GUIDE-seq in HEK293 cells [8], HA-dCas9 in HEK293T cells [12], and POCKET-seq in HCT116 cells for HEK293_s4. CRISPOR predicted 895 sites. GUIDE-seq identified 134 sites. HA-dCas9 identified 484 sites. POCKET-seq identified 3429 sites. Overlaps are quantified between the various techniques.

We also observed little overlap between POCKET-seq and experimental off-target mapping approaches (**Fig. 2D-E**). This difference likely arises from how these assays operate: methods like GUIDE-seq measure direct Cas9 DNA cleavage [8], while DISCOVER-seq+ measures interaction of DNA repair proteins with DSB lesions [9,10]. Because these techniques require cutting to produce a signal, they fail to capture binding-only events. Overall, the larger number of POCKET-seq off-targets compared to other approaches aligns with previous findings that non-cutting interactions between Cas9 nucleases and genomic DNA are more frequent than cutting interactions [11–13].

To account for differences between binding and cleavage assays, we compared POCKET-seq against a previous HA-dCas9 off-target mapping approach [12] (**Fig. 2E**). POCKET-seq identified significantly more off-target sites with little overlap between the two datasets. We attribute this increased sensitivity to our pooled screening approach, which relies on the stable genomic integration and sustained expression of dCas9-KRAB and gRNAs, rather than the transient transfections utilized in previous studies [12]. This difference may also partially reflect cell line-specific chromatin effects.

While each of these techniques samples the underlying landscape of nuclease binding, there are key methodological differences that may influence outcomes. For instance, the pooled CRISPRi screen implementation, described in this manuscript and widely used in the literature, uses dCas9 that is genomically integrated and expressed off a transgenic promoter such as SFFV, while the gRNA is stably expressed from the U6 promoter. These parameters cause moderate but stable levels of RNA-guided catalytically inactive nucleases, which likely drives increased association of these complexes with chromatin. We suggest that dCas9-genome interactions are fundamentally different under these conditions than those used during transient expression experiments. Therefore, characterizing off-targets in the context of transcriptional regulation or epigenome editing experiments will require novel approaches like POCKET-seq, rather than standard approaches developed to model cutting Cas9-genome interactions.

Overall, POCKET-seq provides a highly sensitive method for mapping genome-wide dCas9-KRAB binding events and facilitates interpretation of CRISPRi phenotypes.

## Conclusions

Pooled CRISPRi screens are a powerful approach to identify gene networks that produce a phenotype; however, they are widely known to produce candidate genes that fail to reproduce the original phenotype in secondary analyses. Best practices for pooled screens, such as testing multiple distinct replicates in secondary analyses, prevent false-positive interactions from entering downstream analyses. However, discarding these false positives entirely also discards information that can be used to bolster the primary screen. Here, we developed POCKET-seq to identify off-target interactions in transcriptional regulation experiments.

POCKET-seq reveals off-target binding interactions of dCas9 and classifies these interactions according to their likelihood to produce false-positive or true-positive outcomes. Overall, we believe that the POCKET-seq approach will be an invaluable method to detect relevant off-target events in CRISPRi screens and classify their contributions to screen phenotypes. POCKET-seq is particularly suited for identifying sites of transcriptional repression using dCas9-KRAB that would be missed by alternative approaches. Further development of these methodologies may also be helpful for finding off-targets of genome editors that are based on variants of Cas9, such as prime editors, which utilize nickase Cas9 (nCas9) [14], and base editors, which can utilize dCas9 or nCas9 [15].

## Methods

### Cell Culture

HEK293T cells and HCT116 dCas9-KRAB cells (a gift from the laboratory of J. Corn, ETH Zürich, Zürich, Switzerland) were cultured in Dulbecco’s Modified Eagle Medium (DMEM) GlutaMAX medium (Cat#10569044; Gibco) supplemented with 10% fetal bovine serum (FBS) (Cat#S11550; R&D Systems) and 100 U/mL penicillin/streptomycin (P/S) (Cat#15140122; Gibco). Cell cultures were maintained at 37°C and 5% CO_2_ in a humidified incubator.

### Molecular Cloning

BFP-marked gRNA vectors with puromycin resistance were produced to support CRISPRi. Guides were cloned into the BstXI-BlpI sites. Protospacer sequences can be found in Figure 2A.

### Lentiviral Packaging

Lentivirus was produced by transfecting HEK293T cells with standard delta VPR and VSV-G second-generation packaging vectors. Briefly, approximately 500,000 HEK293T cells were seeded per well in 6-well plates the day before the transfection. The following day, 12 μL of TransIT-LT1 Transfection Reagent (Cat#MIR2305; Mirus Bio) was incubated in 200 μL Opti-MEM Reduced Serum Media (Cat#31985070; Gibco) for 5 min at room temp. Separately, 1.68 μg of delta VPR and 0.42 μg of VSV-G second-generation packaging plasmids were mixed with 2.4 μg of the cargo plasmid (i.e., BFP-marked gRNA vector with puromycin resistance). Next, the plasmids were mixed with the transfection reagents and allowed to incubate for 20 min at room temp. Finally, the solution was added dropwise over the HEK293T cells. At 24H post transfection, the media was changed to remove the transfection complexes. Lentivirus was harvested 72H following transfection, clarified, flash frozen, and stored at −80°C until use.

### Lentiviral Transduction

HCT116 dCas9-KRAB cells were transduced with lentivirus prepared as described above under Lentiviral Packaging. Lentiviral titering was performed to optimize the transduction conditions to ensure single integration events by using a threshold where <20% of the cells were infected. Cells were “spinfected” with lentiviral supernatants by centrifugation at 1,200 x *g* (RCF) for 2H at room temp. Following “spinfection”, the media was changed three times (at 24H, 48H, 72H) to remove lentiviral particles. HCT116 dCas9-KRAB cells expressing BFP-marked gRNA vector were enriched through antibiotic selection using 1.5 μg/mL of puromycin dihydrochloride (Cat#A1113803; Gibco), with drug treatment first starting 48H after lentiviral transduction.

### Lentiviral Addback of SOCS3 cDNA and PEX19 cDNA

Re-expression plasmids were constructed by Gibson Assembly, where each cDNA sequence was inserted into the pLentiX-CD90.1/Thy-1.1 vector backbone. The *PEX19* cDNA addback is denoted as plasmid # pCR2110, while the *SOCS3* cDNA addback is denoted as plasmid # pCR2111. Lentivirus from each of these plasmids was produced as described above under Lentiviral Packaging and transduction was carried out as described above under Lentiviral Transduction. After transducing HCT116 dCas9-KRAB cells with lentivirus, Thy-1.1 positive cells were enriched by Fluorescence-Activated Cell Sorting (FACS). FACS (SH800S Cell Sorter; Sony) was performed by immunolabeling with APC-conjugated anti-CD90.1/Thy-1.1 mouse monoclonal antibody (Cat#17090082; Invitrogen).

### Immunofluorescence Microscopy

Cells were plated on glass-bottom 96-well plates (Cat#P96-1.5H-N; Cellvis) and grown until cell confluency reached between 50% and 80% at the time of fixation. Fixation was performed for 10 min at room temp using a solution of D-PBS containing 4% formaldehyde, which was diluted in house from a concentrated stock of 16% formaldehyde (Cat#28908; Thermo Scientific). After two washes with D-PBS, cells were permeabilized with 0.25% Triton X-100 in D-PBS for 10 min. Next, cells were incubated in blocking buffer (D-PBS containing 0.1% Tween 20 and 3% BSA) for 30 min at room temp. After block, rabbit polyclonal PMP70 primary antibody (Cat#PA1650; Invitrogen) was diluted in blocking buffer and incubated for 1H at room temp with rocking. Cells were then washed three times with PBS-T (D-PBS with 0.1% Tween 20) for 2 min each wash. Secondary antibodies and DAPI (Cat#D1306; Invitrogen) were diluted in blocking buffer and incubated for 1H at room temp in the dark. After incubation, three more washes were performed with PBS-T. Samples were stored in PBS and protected from light prior to image acquisition.

### Peroxisome Quantification by Microscopy

Images were acquired using a Nikon Eclipse Ti2-E inverted microscope configured with a spinning disk confocal scanner (CSU-W1; Yokogawa), an ORCA-Fusion BT sCMOS camera, a CFI Plan Apochromat Lambda D 40X/0.95NA Plan Apo air objective lens, a CFI Apochromat TIRF 100X/1.49NA oil immersion objective lens, and NIS-Elements AR software (v5.31.01; Nikon). Microscopy images were postprocessed using ImageJ/FIJI software (v2.0.0). Quantification and analysis of microscopy images was performed using CellProfiler, Broad Institute (v4.2.4). For live cell images, acquired images were thresholded by global minimum cross entropy to select for and differentiate between cell cytoplasm area and mVenus-PTS foci area in an unbiased manner; downstream mVenus-PTS foci number, area, and intensity was measured within a range of size and ROI. For immunofluorescence microscopy, images were processed by: first, defining nuclei stained by DAPI by adaptive Otsu 3-class thresholding to differentiate between background and nuclei; second, by expanding from nuclei objects to define cytoplasm based on distance and Otsu 2-class thresholding and then subtracting nuclei from this area; third, by selecting, within the cytoplasm area, foci objects for mVenus-PTS or PMP70 of a defined size and ROI determined by adaptive Otsu 3-class thresholding.

### RT-qPCR

Briefly, HCT116 CRISPRi cells harboring target gRNAs were harvested and spun down. RNA was extracted using the RNeasy Mini Kit (Cat#74106; Qiagen) according to the manufacturer’s instructions, in triplicate. RNA was quantified, normalized, and then reverse transcribed using iScript Reverse Transcription Supermix for RT-qPCR (Cat#1708840; Bio-Rad) according to manufacturer’s instructions. The cDNA was then quantified on a CFX96 (Bio-Rad) using 2x SsoFast EvaGreen Supermix (Cat#1725200; Bio-Rad) with corresponding primers for mRNA transcripts for a housekeeping gene (*GAPDH*) and our genes of interest, *SOCS3* and *PEX19*.

### Chromatin Immunoprecipitation

For ChIP-seq, 8 x 10^6^ HCT116 dCas9-KRAB cells were harvested and resuspended in 15 mL of room temp Dulbecco’s Modified Eagle Medium (DMEM) GlutaMAX medium (Cat#10569044; Gibco) without serum and without antibiotics. Cells were crosslinked for 10 min at room temp in suspension with rotation by the addition of 1 mL of 16% formaldehyde (Cat#28908; Thermo Scientific) (final formaldehyde concentration of 1%). The reaction was quenched for 5 min at room temp in suspension with rotation by the addition of 0.825 mL of 2.5 M glycine (final glycine concentration of 125 mM). After three washes with ice-cold D-PBS, centrifuging at 1,200 x *g* (RCF) for 3 minutes each wash, the cells were flash frozen and stored at −80°C for future processing. Cell lysis and nuclear fractionation were carried out using a series of chilled lysis buffers. First, the cell pellets were resuspended in 1 mL LB1 lysis buffer (50 mM HEPES–KOH, pH 7.5; 140 mM NaCl; 1 mM EDTA; 10% Glycerol; 0.5% IGEPAL CA-630; 0.25% Triton X-100) containing protease inhibitors (Cat#PI78429; Fisher Scientific) and incubated for 10 min at 4°C with rotation. Next, the samples were then centrifuged at 2,000 x *g* (RCF) for 3 min, and the new pellets were resuspended in 1 mL LB2 lysis buffer (10 mM Tris–HCl, pH 8.0; 200 mM NaCl; 1 mM EDTA; 0.5 mM EGTA) containing protease inhibitors (Cat#PI78429; Fisher Scientific) and incubated for 5 min at 4°C with rotation. The samples were again centrifuged at 2,000 x *g* (RCF) for 3 minutes, and the new pellets were resuspended in 0.5 mL LB3 lysis buffer (10 mM Tris– HCl, pH 8.0; 100 mM NaCl; 1 mM EDTA; 0.5 mM EGTA; 0.1% Na–Deoxycholate; 0.5% N-Lauroylsarcosine) containing protease inhibitors (Cat#PI78429; Fisher Scientific). Chromatin was sheared in LB3 lysis buffer by sonication using a Qsonica Q500 sonicator with a Cup Horn, until the average shear size ranged from 200-600 bp (24 cycles of 15 sec on, 45 sec off, at 50% amplitude, for a total on time of 6 min). Sonicated lysates were cleared by centrifugation at 21,100 x *g* (RCF) at 4°C for 10 min. Immunoprecipitations were carried out using the cleared, sonicated lysate, which was brought up to 1 mL final volume with additional LB3 lysis buffer and 0.1 mL of 10% Triton X-100 (final Triton X-100 concentration of 1%). After saving 5% of each sample as input, each IP sample received 50 μL of Dynabeads protein A magnetic beads (Cat#10001D; Invitrogen) that were pre-conjugated to 5 μg of anti-HA tag ChIP-grade antibody (Cat#ab9110; Abcam). Immunoprecipitations were carried out overnight at 4°C with rotation. Samples were then washed 5x with ice-cold ChIP wash buffer (50 mM HEPES–KOH, pH 7.5; 500 mM LiCl; 1 mM EDTA; 1% IGEPAL CA-630; 0.7% Na–Deoxycholate) for 5 min at 4°C with rotation per wash. Samples were then washed 1x with ice-cold TBS (20 mM Tris–HCl; 150 mM NaCl) for 5 min at 4°C with rotation. Samples were eluted from the magnetic beads using two 15-min incubations in ChIP elution buffer (0.1 M NaHCO_3_, 1% SDS) at 65°C on a heated shaker at 1200 rpm. Input and IP samples (5%) were collected for Western Blotting to confirm enrichment (described below). Crosslinks were reversed overnight at 65°C on a heated shaker at 1200 rpm using 200 mM final concentration NaCl. Samples were purified using the MinElute PCR Purification Kit (Qiagen #28004) after RNase and Proteinase K treatment.

### High Throughput Sequencing

DNA library preparation of the ChIP samples was commercially performed (Novogene, Beijing). Next Generation Sequencing (NGS) was performed using the NovaSeq PE150 (Illumina) platform (Novogene, Beijing).

### Western Blot

Input and IP samples from ChIP experiments were mixed 1:1 with 2X Laemmli blue sample buffer (20% glycerol; 125 mM Tris–HCl, pH 6.8; 4% SDS; 5% beta-mercaptoethanol; 0.02% bromophenol blue). Samples were boiled in sample buffer for 5 minutes at 95°C. Samples were separated via electrophoresis on 4–20% TGX stain-free SDS-PAGE gels (Bio-Rad #4568093). Before transfer, the TGX stain-free chemistry of the SDS-PAGE gels was activated for 45 seconds and subsequently used as a loading control. Gels were transferred using a semi-dry apparatus onto 0.45 µm low fluorescence PVDF membranes (Bio-Rad #1620264). After transfer, membranes were blocked for 1 hour in 5% non-fat dry milk in TBS (20 mM Tris–HCl, pH 7.5; 150 mM NaCl) with 0.05% Tween-20. Membranes were incubated overnight at 4°C in anti-HA tag mouse monoclonal HA.C5 antibody (Abcam # ab18181) with a dilution of 1:1,000. Membranes were washed with TBS with 0.05% Tween-20, and then incubated for 1 hour at room temp with goat anti-mouse IgG (H + L)-HRP conjugate antibody (Bio-Rad # 1706516) with a dilution of 1:3,000. Pierce ECL 2 Western Blotting Substrate (Thermo Scientific # PI80196) was used to visualize membranes by chemiluminescence (Bio-Rad ChemiDoc MP Imaging System).

### Generation of pooled screen phenotype scores

NGS data was quantified, and phenotype scores were generated using python scripts from the Horlbeck Lab’s Screen Processing pipeline as previously described [16] and **MAGeCK-iNC** [4,5].

### POCKET-seq Data Analysis

FASTQ files generated by high-throughput sequencing were aligned to the human reference genome (hg19) using Bowtie 2 [17]. Aligned reads were processed with SAMtools [18] to produce sorted, indexed BAM files. Peak calling was performed on the resulting BAM files using MACS3 [19] in paired-end mode to identify genomic regions enriched for dCas9-KRAB binding. MACS3 output was annotated using a custom python script (POCKET-seq, available at https://github.com/RichardsonDNARepair/POCKET-seq). POCKET-seq is a python3 script requiring packages pandas [20], NumPy [21], and argparse. For each peak, POCKET-seq calculates the peak midpoint and identifies the nearest transcript in a provided GFF3 annotation file using the TSS as the reference coordinate in a strand-aware manner. Each peak is annotated with the nearest gene symbol, its distance in kilobases (kb) from the peak midpoint to the TSS, and a binary flag indicating whether that distance falls within 1 kb. Where primary screen phenotype data are available, peaks can additionally be annotated with the epsilon score of the nearest gene. For gene ontology analysis, peaks are flagged based on membership of the nearest gene in a user-specified GO term using an index built from GO annotation (GAF) and ontology (OBO) files downloaded from geneontology.org [22,23].

## Declarations

### Ethics approval and consent to participate

Not applicable.

### Consent for publication

Not applicable.

### Availability of data and materials

The datasets generated and/or analyzed during the current study are available in the SRA repository under accession number PRJNA1473212. Primary screen data is used to annotate POCKET-seq results. This screen is published in PMID: 38967608 [6] and the raw data is available through GEO (GSE266855).

### Authors’ contributions

C.D.R., B.M.G., J.T.V., and C.M.J. conceived of the POCKET-seq method. J.T.V. and K.U.T. developed cell lines and reagents. C.M.J., J.T.V., and G.D.K. collected data. G.D.K. and C.D.R. analyzed data. C.D.R. and B.M.G. supervised the project. C.M.J., G.D.K., and C.D.R. wrote the manuscript with input from all authors. All authors read and approved the final manuscript.

## Acknowledgements

We thank Katelynn Kazane for assistance with engineering CRISPRi cell lines. We thank Amos Liang (University of Dundee, Dundee, Scotland, UK) and Jacob Corn (ETH Zurich, Zurich, Switzerland) for HCT116 CRISPRi cells. We thank Mary West and Pingping He of the High-Throughput Screening Facility at the University of California, Berkeley for preparation of the pooled CRISPRi library and Jonathan Weissman (Whitehead Institute for Biomedical Research, Boston, MA, USA) for the gift of the pooled library. We thank members of the Gardner and Richardson labs for their feedback on the manuscript. The authors acknowledge the assistance of Dr. Jennifer Smith, manager of the Biological Nanostructures Laboratory within the California NanoSystems Institute, supported by the University of California, Santa Barbara and the University of California, Office of the President.

## Supplementary Information

**Supplemental Figure 1.**
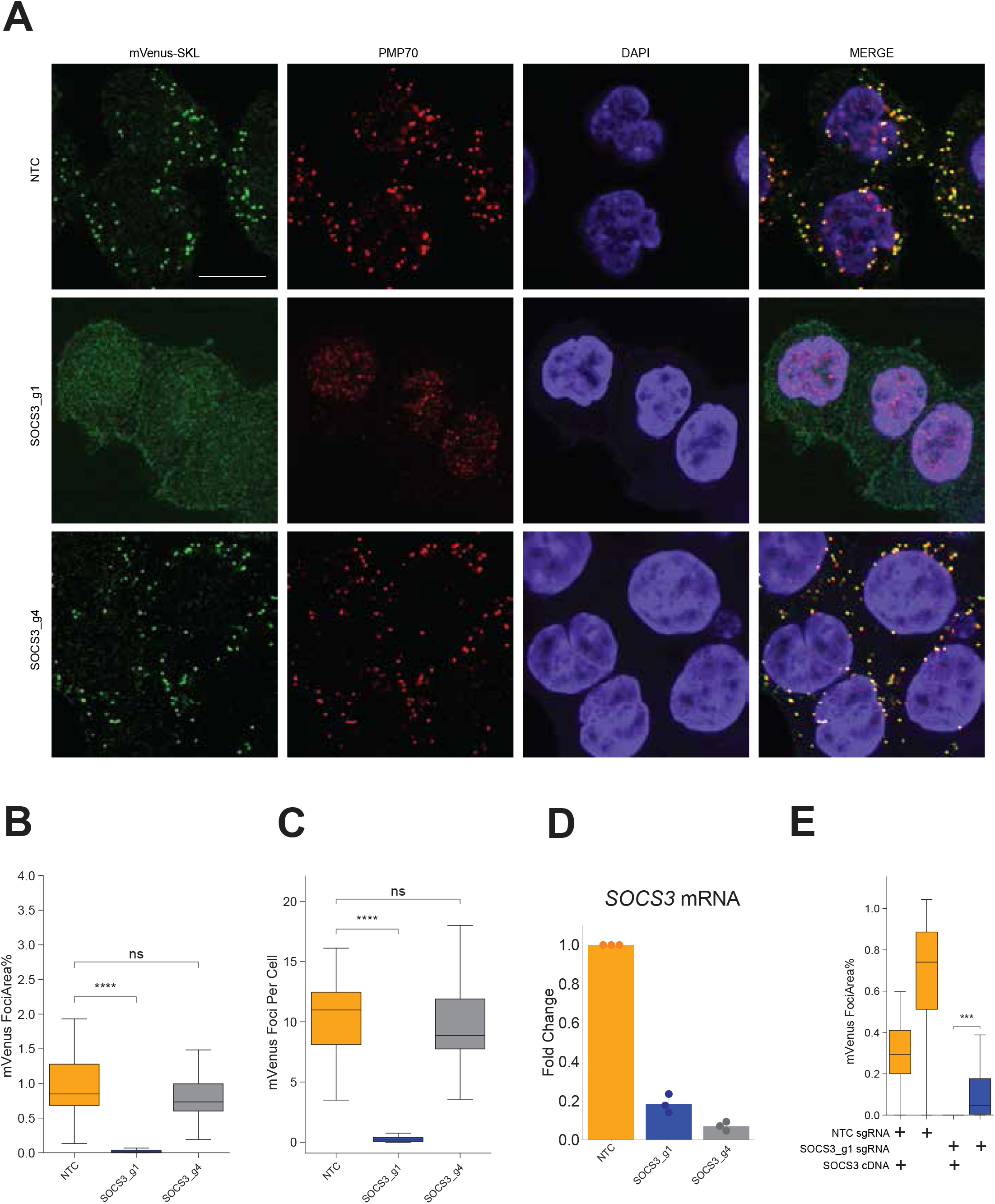
A. Representative microscopy images showing mVenus-PTS (fluorescent reporter) and PMP70 (immunofluorescence staining) in CRISPRi non-targeting control (NTC) and *SOCS3*-depleted (SOCS3_g1 and SOCS3_g4) HCT116 cells. Data shown are generated from at least *m* = 24 images and *n* = 500 cells. Scale bar represents 10 µm. B. Quantification of mVenus-PTS foci area from panel S1A as a percentage of cell area. The *p* values were derived from an independent t-test. *ns*, not significant; \*\*\*\**p* ≤ 0.0001. C. Quantification of mVenus-PTS foci per cell from panel S1A. The *p* values were derived from an independent t-test. *ns*, not significant; \*\*\*\**p* ≤ 0.0001. D. RT-qPCR for *SOCS3* mRNA levels in CRISPRi non-targeting control (NTC) and *SOCS3*-depleted (SOCS3_g1 and SOCS3_g4) HCT116 cells. Data represented as the fold change of *SOCS3* transcript levels normalized to NTC. Data shown are representative of *n* = 3 biological replicates. E. Quantification of mVenus-PTS foci area as a percentage of cell area from live cell microscopy of mVenus-PTS (fluorescent reporter) in CRISPRi non-targeting control (NTC) and *SOCS3*-depleted (SOCS3_g1) HCT116 cells with either no addback or addback of *SOCS3* cDNA. Addbacks are stably expressed under hEF1a promoters. Data shown are generated from at least *m* = 32 images and *n* = 1500 cells. The *p* values were derived from an independent t-test. \*\*\**p* ≤ 0.001. Representative microscopy images can be found in S2C.

**Supplemental Figure 2.**
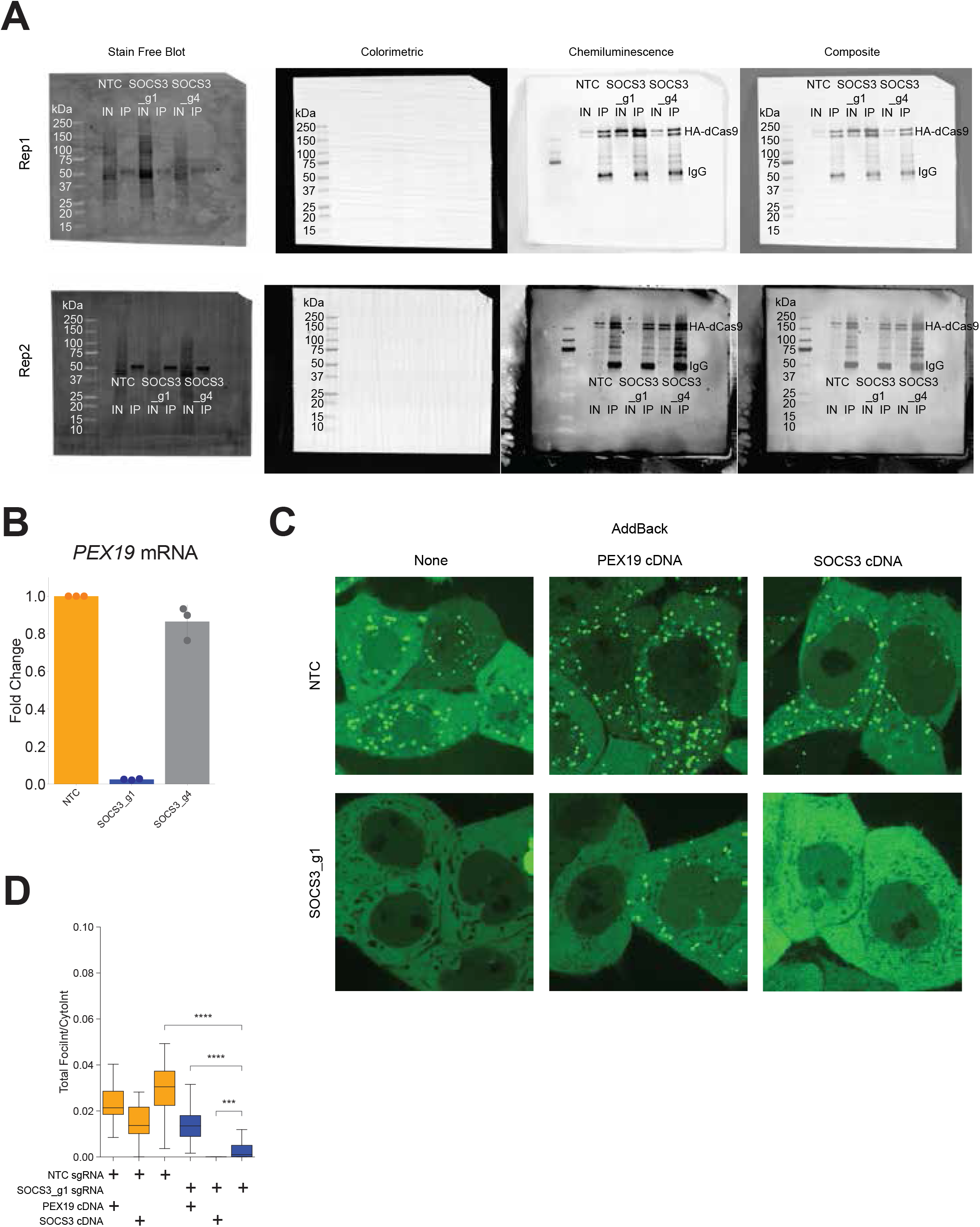
A. Raw western blots of input (IN) and anti-HA tag immunoprecipitation (IP) fractions during the POCKET-seq ChIP procedure from CRISPRi non-targeting control (NTC) and *SOCS3*-depleted (SOCS3_g1 and SOCS3_g4) HCT116 cells where dCas9-KRAB is epitope tagged with HA. From left to right, blots show total protein (Stain Free), protein ladder (Colorimetric), anti-HA tag signal (Chemiluminescence), and merge (Composite). B. RT-qPCR for *PEX19* mRNA levels in CRISPRi non-targeting control (NTC) and *SOCS3*-depleted (SOCS3_g1 and SOCS3_g4) HCT116 cells. Data represented as the fold change of *PEX19* transcript levels normalized to NTC. Data shown are representative of *n* = 3 biological replicates. C. Representative live cell microscopy images showing mVenus-PTS (fluorescent reporter) in CRISPRi non-targeting control (NTC) and *SOCS3*-depleted (SOCS3_g1) HCT116 cells with either no addback, addback of *PEX19* cDNA, or addback of *SOCS3* cDNA. Addbacks are stably expressed under hEF1a promoters. Data shown are generated from at least *m* = 32 images and *n* = 1500 cells. F. Quantification of total mVenus-PTS foci intensity over total cytoplasmic mVenus-PTS intensity from panel S2C. The *p* values were derived from an independent t-test. \*\*\**p* ≤ 0.001; \*\*\*\**p* ≤ 0.0001.

